# Multiple loci control eyespot number variation on the hindwings of *Bicyclus anynana* butterflies

**DOI:** 10.1101/653451

**Authors:** Angel G. Rivera-Colón, Erica L. Westerman, Steven M. Van Belleghem, Antónia Monteiro, Riccardo Papa

## Abstract

The underlying genetic changes that regulate the appearance and disappearance of repeated traits, or serial homologs, remain poorly understood. One hypothesis is that variation in genomic regions flanking master regulatory genes, also known as input-output genes, controls variation in trait number, making the locus of evolution almost predictable. Other hypotheses implicate genetic variation in upstream or downstream loci of master control genes. Here, we use the butterfly *Bicyclus anynana*, a species which exhibits natural variation in eyespot number on the dorsal hindwing, to test these two hypotheses. We first estimated the heritability of dorsal hindwing eyespot number by breeding multiple butterfly families differing in eyespot number, and regressing eyespot number of offspring on mid-parent values. We then estimated the number and identity of independent genetic loci contributing to eyespot number variation by performing a genome-wide association study with restriction site-associated DNA Sequencing (RAD-seq) from multiple individuals varying in number of eyespots sampled across a freely breeding lab population. We found that dorsal hindwing eyespot number has a moderately high heritability of approximately 0.50. In addition, multiple loci near previously identified genes involved in eyespot development display high association with dorsal hindwing eyespot number, suggesting that homolog number variation is likely determined by regulatory changes at multiple loci that build the trait and not by variation at single master regulators or input-output genes.

**Data accessibility:** The *Bicyclus anynana* PstI RAD-tag sequencing data is available via the Genbank BioProject PRJNA509697. Genotype VCF files will be made available through Figshare upon acceptance.

## Introduction

Body plans often evolve through changes in the number of repeated parts or serial homologs by either addition or subtraction. For instance, the pelvic fins of vertebrates are inferred to have originated by addition to a body plan displaying only pectoral fins, perhaps via co-option of the pectoral or caudal fin developmental programs to a novel location in the body (Ruvinsky and Gibson-Brown 2000; Larouche *et al.* 2017). In insects, the absence of limbs and wings in the abdomen is inferred to be due to a process of subtraction, i.e. via the repression or modification of limbs and wings in these segments by *hox* genes (Galant and Carroll 2002; Ronshaugen *et al.* 2002; Tomoyasu *et al.* 2005; Ohde *et al.* 2013). Regulatory targets of abdominal *hox* genes are likely to underlie loss of limb/wing number in these body segments (Tomoyasu *et al.* 2005; Ohde *et al.* 2013), although these mutations have not yet been identified. Thus, while serial homolog number variation is a common feature in the evolution of organisms’ body plans, the underlying genetic changes that regulate the appearance and disappearance of these repeated traits remain poorly understood.

Studies in *Drosophila* have contributed most to the identification of the genetic basis underlying the evolution of serial homolog number. Larvae of different species have different numbers of small hairs, or trichomes, in their bodies, and variation in regulatory DNA around the gene *shavenbaby* appears to be largely responsible for this variation (McGregor *et al.* 2007). Moreover, *shavenbaby* has been labeled a master regulatory gene because its ectopic expression in bare regions of the body leads to trichomes (Payre *et al.* 1999). However, a more complex genetic architecture seems to underlie variation in the number of larger bristles found in the thorax of adults. In this case variation around *achaete-scute*, a gene complex required for bristle differentiation, plays a role in controlling bristle number variation across species (Marcellini and Simpson 2006). Interestingly, genetic variation in upstream regulatory factors, whose spatial expression overlaps some, but not all bristles, is also known to impact bristle number in lab mutants (Garcia-Bellido and de Celis 2009). Finally, *shavenbaby* and *scute* genes are also known as input-output genes due to their central “middle of the hour-glass” position in regulatory networks (Stern and Orgogozo 2008). These genes respond to the input of multiple upstream protein signals, present at distinct locations in the body, and in turn control the regulation of the same battery of downstream genes, to affect the same output (trichome or bristle development) at each of these body locations. Mutations in the regulatory regions of these genes are thus expected to have minimal pleiotropic effects, and to lead to changes in the number of times the network is deployed, and thus to evolution in the number of trichome or bristles in the bodies of these flies. While this type of regulatory network architecture points to predictable regions in the genome that will readily evolve leading to trait number evolution, i.e., hotspots of evolution, it might represent only one type of architecture among others that are still unexplored. More systems, thus, need to be investigated for a more thorough understanding of the genetic basis underlying variation of repeated traits in bodies.

One promising system for investigating the genetic basis of serial homolog number evolution is the group of eyespot patterns on the wings of nymphalid butterflies. Eyespots originally appeared on the ventral hindwing in a lineage of nymphalid butterflies, sister to the Danainae, and have subsequently been added to the forewings and dorsal surfaces of both wings (Oliver *et al.* 2012, 2014; Schachat *et al.* 2015). Furthermore, within a single species, eyespot number can vary significantly between individuals or sexes (Brakefield and van Noordwijk 1985; Owen 1993; Tokita *et al.* 2013), allowing for population genetic approaches to identify the underlying genetic basis of such variation. Genes controlling eyespot number variation within a species might also be involved in promoting this type of variation across species.

One of the best model species for studying the genetic basis of eyespot number variation is the nymphalid butterfly *Bicyclus anynana*. This species exhibits natural variation and sexual dimorphism in eyespot number on the dorsal hindwing surface, which plays a possible role in mate choice (Westerman *et al.* 2014). The observed variation consists of males averaging 0.75 dorsal hindwing eyespots, with a range of 0-3, and females averaging 1.5 dorsal hindwing eyespots, with a range of 0-5 (Westerman *et al.* 2014) (Fig. 1). Dorsal hindwing eyespot number variation is positively correlated with butterfly size, and, in general, spots are added sequentially on the wing (Cu1, M3, then either Cu2 or M2, and, rarely, Pc). Lab populations of this species also display a series of mutant variants that affect eyespot number on other wing surfaces. Genetic and developmental studies on eyespot number variation in this species suggest the existence of at least two different underlying molecular mechanisms. Spontaneous mutants such as Spotty (Brakefield and French 1993; Monteiro *et al.* 1997, 2013), Missing (Monteiro *et al.* 2007), P- and A- (Beldade *et al.* 2008), or X-ray induced mutations such as 3+4 (Monteiro *et al.* 2003), segregate as single Mendelian alleles, and cause discrete and obvious changes in eyespot number or affect the size of very specific eyespots. On the other hand, multiple alleles of small effect likely regulate the presence or absence of small eyespots that sometimes appear between the typical two eyespots on the forewing, or on the most posterior wing sector of the ventral hindwing. This type of eyespot number variation is positively correlated with eyespot size variation, responds readily to artificial selection on eyespot size (Holloway *et al.* 1993; Monteiro *et al.* 1994; Beldade and Brakefield 2003), and is likely under the regulation of a threshold type mechanism (Brakefield and van Noordwijk 1985).

**Figure 1.**
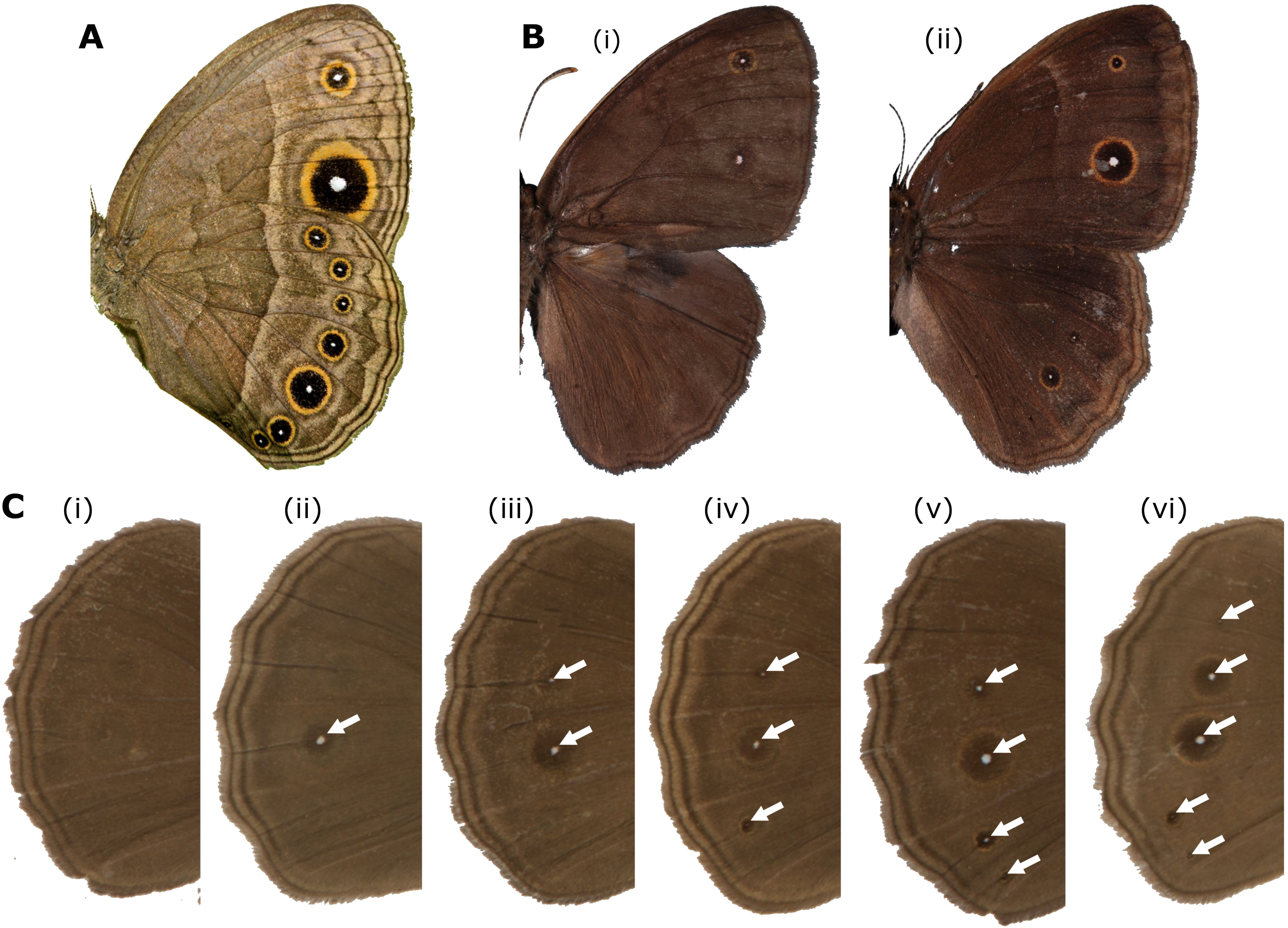
Eyespot pattern and number variation in *Bicyclus anynana*. **(A.)** Eyespot pattern on the ventral side of wings: forewing displays two eyespots; hindwing displays seven eyespots. **(B.)** Eyespots pattern on dorsal side of wings. Male (left) displaying two dorsal forewing eyespots and zero dorsal hindwing eyespots and female displaying two dorsal forewing eyespots and three dorsal hindwing eyespots. **(C.)** Dorsal hindwing eyespot number (DHEN) variation, ranging from zero to five UV-reflective spots, marked by white arrows (i–vi).

Interestingly, eyespot number variation within *B. anynana* can involve changes to single eyespots or to several eyespots at a time, on one or both wing surfaces. For instance, Spotty introduces two eyespots on the dorsal and ventral surfaces of the forewing, whereas A- and P- primarily reduce the size of the single anterior (A-) or the posterior eyespot (P-) of the dorsal surface exclusively, without affecting eyespot size or number on the ventral surface. The genetic basis for these differences is still unknown.

Recently, the gene *apterousA (apA)* was shown to regulate wing pattern differences between dorsal and ventral surfaces in *B. anynana*, including differences in eyespot number (Prakash and Monteiro 2018). This gene is expressed exclusively on the dorsal wing surfaces and its mutation via CRISPR-Cas9 led to dorsal wing surfaces acquiring a ventral identity, which included additional eyespots. This study indicated that *apA* is a repressor of eyespots on the dorsal surface. However, *B. anynana*, has eyespots on dorsal wing surfaces and their presence and variation in number appears to be correlated with variation in the number of small circular patches, positioned at future eyespot centers, lacking *apA* expression (Prakash and Monteiro 2018). This suggests that genetic variation at loci that modulate the expression of *apA* in eyespot centers on the dorsal surface, or genetic variation in regulatory regions of *apA* itself, might be involved in regulating eyespot number specifically on the dorsal surface of wings.

The genetic architecture of eyespot number variation in any butterfly species remains unknown. Here, we examine the genetic basis of dorsal hindwing eyespot number (DHEN) variation in *B. anynana*. We carried out two sets of experiments. We first estimated the heritability for this trait by breeding multiple butterfly families differing in eyespot number and regressing eyespot number of offspring on mid-parent values. Then we estimated the number and identity of independent genetic loci that are contributing to variation in this trait by performing a genome-wide association study with restriction site-associated DNA Sequencing (RAD-seq) from multiple individuals varying in number of eyespots sampled across a freely breeding lab population.

## Materials and Methods

### Study organism

*Bicyclus anynana* is a Nymphalid butterfly common to sub-tropical Africa for which a colony has been maintained in the laboratory since 1988. All *Bicyclus anynana* butterflies used in this study were collected from a colony established in New Haven, CT (Yale University), composed of an admixed population of numerous generations of freely breeding individuals with variable dorsal hindwing eyespot number phenotypes. Individuals from this colony originated from an artificial colony established in Leiden University, from numerous gravid females collected in Malawi in 1988. These laboratory populations have been maintained at relatively high populations sizes, and females preferentially avoid mating with inbred males (van Bergen *et al.* 2013), which facilitates the relatively high levels of heterozygosity detected several years post establishment (Saccheri and Bruford 1993; Saccheri *et al.* 1996). The colony was kept in controlled conditions of 12 hours light/dark cycles, 80% relative humidity and a temperature of 27°C. Larvae were fed on corn plants and adult butterflies on mashed banana, as described in previous publications (Westerman *et al.* 2014).

### Heritability of dorsal hindwing eyespot number

We examined the number of dorsal hindwing eyespots (DHEN) on all offspring from 18 separately reared families whose parents differed in eyespot number: six families where both parents had DHEN of zero (0F x 0M); six where both parents had DHEN of one (1F x 1M); and six where both parents had DHEN of two (2F x 2M). All generations were reared in the conditions described above. We ensured virginity of the females by separating the butterflies in the parental generation into sex-specific cages on the day of eclosion. All families were started within 5 days of each other using adults ranging from 1-3 days old. While initial adult age ranged from 1-3 days, the average age was consistent across the three DHEN treatments (ANOVA, n=18, DF=2, F=0.8266, p=0.4565). Each breeding pair was placed in a cylindrical hanging net cage of 30 cm diameter X 40 cm height, with food (banana slices), water and a young corn plant on which to lay eggs. When corn plants were covered with eggs, they were placed in family-specific mesh sleeve cages for larval growth. Females were given new plants on which to lay eggs until they died. Pupae and pre-pupae were removed from the sleeve cages and placed in family-specific cylindrical hanging net cages for eclosion. The cages were checked daily for newly emerged butterflies. On the day of eclosion, DHEN was recorded for each offspring. DHEN was calculated for each individual by averaging the number of dorsal hindwing eyespots on the left and right wing, which allowed for intermediate values when the wings were asymmetric. Narrow-sense heritability (*h*^2^) was calculated by regressing offspring on midparent values and correcting the heritability estimate for assortative mating. This corrected heritability value 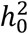 was calculated using correlations between phenotypic values of mated pairs (*r*) to calculate the correlation between breeding values of mates (m = *rh*^2^), with 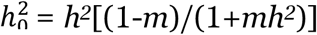 as described in Falconer & Mackay (1996). Estimates were obtained for the pooled offspring data as well as for separate regressions of female and male offspring data on mid-parent values. Sex-specific heritabilities were calculated using the correction for unequal variances in the two sexes (for example, regression of daughter on father, adjusted regression *h*’ = *b*σ_m_/σ_f_ with σ_m_/σ_f_ the ratio of phenotypic standard deviations of males to females) (Falconer and Mackay 1996). Given the known effect of sex on DHEN (Westerman *et al.* 2014), we then also tested for an interaction of parental phenotype and offspring sex on offspring phenotype using a general linear model, with sex, parental phenotype, and sex*parental phenotype as fixed variables.

### Sample collection and phenotype determination for genomic association study

To identify regions in the genome that are associated with DHEN variation, we collected and sequenced a total of 30 individuals from the previously described laboratory population. Fifteen individuals contained no eyespots (absence) and 15 containing two or more eyespots (presence) (Table S1). Both groups contained an assorted number of male and female individuals. Wings of the collected individuals were removed, and the bodies were preserved in ethanol for DNA extraction.

### RAD library preparation and sequencing

Genomic DNA of the preserved bodies was extracted using DNeasy Blood & Tissue Kit (Qiagen), with an additional RNase digestion for removing RNA from the extracted nucleic acid samples. The quality and concentration of the extracted DNA was verified using gel electrophoresis and Qubit 2.0 fluorometer (Life Technologies). Extracted genomic DNA was used for preparing Illumina RAD sequencing libraries based on previously described protocols (Baird *et al.* 2008; Etter *et al.* 2011). DNA was digested with the frequent cutting enzyme *PstI* and ligated to P1 adapters containing a unique barcode 5 bp in length. Samples were pooled and sheared using a Covaris M220 (Covaris Inc.) instrument and size selected for 300-500 bp inserts on average. After end-repair and P2 adapter ligation, the library was amplified by PCR. The pooled library was then sequenced utilizing a single lane of an Illumina HiSeq2000® 100 bp paired-end module.

### Read quality and filtering

Following sequencing of a RAD-seq library composed of 30 *B. anynana* individuals, we obtained 127 million paired-end reads 100 bp in length. The raw RAD reads were demultiplexed using *Stacks v1.42* (Catchen, Hohenlohe, Bassham, Amores, & Cresko, 2013; Catchen *et al.*, 2011) *process_radtags* pipeline, and reads with low quality and/or ambiguous barcodes were discarded. Further, we removed Illumina adapter sequences from the reads and trimmed sequences to 80 bp in length, as suggested from the FastQC quality control tool (http://www.bioinformatics.babraham.ac.uk/projects/fastqc/) results. We retained a total of 111 million (86 %) and an individual average of 3.7 million ± 1.2 million filtered paired-end reads.

### Reference alignment

The 111 million retained read pairs were aligned to the most recent *B. anynana* genome assembly (v1.2)shiv with its corresponding annotation (Nowell *et al.* 2017). This reference assembly is highly fragmented and composed of 10,800 scaffolds with an N50 of 638.3 Kb, a total assembly size of 475 Mb, and 22,642 annotated genes. Filtered reads were aligned to the reference genome using BWA v0.7.13 (Li and Durbin 2009) *mem* with default seed lengths, mismatch and gap scores, but allowing for the marking of shorter split reads as secondary alignments for compatibility with PicardTools v1.123 (https://broadinstitute.github.io/picard/). Resulting alignments were directly converted to BAM files using SamTools v1.12 (Li *et al.* 2009a) *view*. BAM files were then sorted with SamTools *index* and filtered for duplicates using PicardTools v1.123 *MarkDuplicates* and processed with *AddOrReplaceReadGroups* for GATK compatibility. In total, we obtained a 92.9 % read alignment and 79.6 % properly mapped read pairs. At each RAD locus, average per-individual sequencing coverage was 18.8X (± 4.3; median: 18.3).

### Variant calling and association mapping

After aligning reads to the reference genome, we used two parallel analytical procedures to guarantee our results were method agnostic. First, the processed and filtered alignments were genotyped using GATK v3.5 (McKenna *et al.* 2010) *UnifiedGenotyper* with the default call confidence values and only outputting variant sites. The resulting 4,661,849 raw variant sites were then filtered using VCFTools v0.1.14 (Danecek *et al.* 2011) *recode* to obtain only calls with a minimum genotype quality of 30, a minimum genotype depth of 5, present in over 50 % of all individuals, a max allele number of 2, and minimum allele frequency of 0.05. This results in a filtered VCF containing only 350,121 high-quality biallelic variants.

In parallel, the filtered mapped reads were genotyped using *Stacks* v1.42 (Catchen *et al.* 2011, 2013). Using the *ref_map.pl* wrapper script with default parameters, RAD loci were assembled from the mapped reads using *pstacks*. A catalog of all loci was generated with *cstacks* and samples were matched to this catalog using *sstacks*. The *populations* program was then run on this catalog to generate population genetic measures, enabling the calculation of *F_ST_* statistics. For a variant to be included in the analysis, it had to be present in both study groups (p=2) in over 75% percent of individuals (r=0.75) and have a minimum allele frequency above 0.05. Using *Stacks* we reconstructed 207,752 RAD loci and 673,340 raw SNPs. After filtering, 73,159 RAD loci and 238,786 SNPs were retained. Filtered variants obtained from both datasets were compared to retain only SNPs with support from both genotyping methods. A total of 216,338 SNPs were shared between the two datasets and were used for subsequent comparisons.

To identify areas of the genome associated with our desired phenotype, we scanned the genome using two methods: *F_ST_* and association via a relatedness-corrected linear mixed model. *F_ST_* for all 216 thousand shared variants was calculated using the Stacks *populations* module. A Fisher’s Exact Test *p*-value correction was applied to the resulting *F_ST_* and Analysis of Molecular Variance (AMOVA) *F_ST_* values as a multiple testing correction. Future references to *F_ST_* in the manuscript specifically refer to the corrected AMOVA *F_ST_* values from this analysis. During this *Stacks* run, we enabled the --*plink* flag to output our 216 thousand variant sites into *Plink*’s .*map* and .*ped* format for use in the association analysis. The generated .*map* and .*ped* files were converted to binary .*bed* files using *Plink* v1.90b6.9 (Purcell *et al.* 2007; Clarke *et al.* 2011), in addition to adding phenotype information to all samples. From this, we generated a relatedness matrix of all individuals using *GEMMA* v0.98.1 (Zhou and Stephens 2012, 2014), which was used as one of the parameters of a *GEMMA* univariate linear mixed model. We used the *p*-values of the likelihood ratio tests as the basis of the phenotype to genotype association.

### Selection of candidate loci

To minimize the identification of false genotype-to-phenotype relationships, only areas of the genome displaying both association and *F_ST_* outliers were used for further analysis. The use of both metrics simultaneously ensured that the relationships observed are method agnostic. This multi-metric approach, including a combination of association and *F_ST_* outliers, has been utilized repeatedly to identify genomic regions associated with domestication in both dogs and cats (Axelsson *et al.* 2013; Montague *et al.* 2014), variation in feather coloration in warblers (Brelsford *et al.* 2017), and the architecture and modularity of wing pattern variation in *Heliconius* butterflies (Nadeau *et al.* 2014; Van Belleghem *et al.* 2017). The relationship between the two metrics was explored by plotting both a correlation and a quantile-quantile plot (Q-Q plot) of the *F_ST_* and association for all variants (Fig. S1). Significant peaks were defined as areas of the genome with SNPs containing association and *F_ST_* 6 standard deviations above the genome-wide mean. A second threshold was established for variants with *p*-value association and *F_ST_* at least 7 standard deviations above the mean. We explored all annotated genes within a 500 kb window centered around outlier SNPs, and especially under the strongest associated peak area. To identify direct effects of the outlier SNPs over the genes within these windows, we annotated the variants using *SnpEff* v4.3T (Cingolani *et al.* 2012), building a *de novo* database with the available *B. anynana* reference annotation.

### Principal Component Analysis

To determine the baseline-level genome-wide diversity and divergence among individuals in the present population, we performed a Principal Component Analysis (PCA) on the obtained genotypes. Although our sampled individuals originated from a single, freely breeding population, performing this analysis corroborates that the observed genomic diversity lacks potential substructuring that could impact our outlier identification. To do this, we randomly selected a subset of 5000 filtered variants from the *Stacks* catalog and made them into a whitelist, as described by the *Stacks* manual and by Rochette & Catchen (2017). We then ran the *populations* module on this subset of variants with the addition of the --*genepop* export format flag. The resulting genepop file was processed using the *adegenet* v2.1.1 R package (Jombart 2008; Jombart and Ahmed 2011) by converting the genotype calls into a *genind* object, scaling missing data by mean allele frequency, and analyzed with PCA.

### Ordering of the *B. anynana* scaffolds along the Heliconius melpomene genome

The current *B. anynana* reference assembly (Nowell *et al.*, 2017) has an N50 of 638.3 kb and is composed of 10,800 unlinked scaffolds. To assess whether associated SNPs on separate *B. anynana* genome scaffolds could be part of the same block of association, we ordered the scaffolds of the *B. anynana* genome along the *Heliconius melpomene* v2 genome assembly (Davey *et al.* 2016). Although *B. anynana* and *H. melpomene* diverged about 80 My ago (Espeland *et al.* 2018) and have a different karyotype (n=28 in *B. anynana* versus n=21 in *H. melpomene*), the *H. melpomene* genome is the most closely related butterfly genome that has been assembled into highly contiguous chromosomal scaffolds using pedigree informed linkage maps. Aligning both genomes provides valuable information to interpret our association analysis. To construct this alignment, we used the alignment tool *promer* from the MUMmer v3.0 software suite (Kurtz *et al.* 2004). *Promer* was used with default settings to search for matches between sequences translated in all six possible reading frames between the *B. anynana* and *H. melpomene* genome. The obtained alignments were subsequently filtered for a minimum alignment length of 200 bp and a minimum percent identity (%IDY = (matches x 100)/(length of aligned region)) of 90 %. These filtered alignments were used to order the *B. anynana* scaffolds according to the order in which they aligned along the *H. melpomene* genome. If a scaffold aligned to multiple locations or chromosomes, priority was given to the position it matched with highest identity. For scaffolds that contained significant associations with hindwing eyespot number, we also retained alignments with a minimum %IDY of 70 % and a minimum alignment length of 150 bp to investigate possible fine scale rearrangements between the *B. anynana* and *H. melpomene* genome.

### Linkage disequilibrium analysis

In addition to ordering the *B. anynana* scaffolds to the *H. melpomene* genome for assessing the genomic linkage of SNPs, we calculated linkage disequilibrium in our *B. anynana* study population. To calculate linkage disequilibrium for genomic SNPs, we phased 213,000 SNPs that were genotyped in all samples using *beagle* v4.1 (Browning and Browning 2007). Estimates of linkage disequilibrium were calculated from 100,000 randomly selected SNPs, using the *VCFtools* v0.1.14 (Danecek *et al.* 2011) –*hap-r2* function, with a max LD window of 5 Mbp, and minimum allele frequency cutoff of 0.10. Resulting LD comparisons for genomic SNPs were then plotted in R, where a Loess local regression was calculated and used to determine the genome-wide window size of linkage disequilibrium decay. Subsequently, this LD window size was used for the investigation of genes near associated loci.

## Results

### Dorsal hindwing spot number variation has moderate to high heritability

Zero spot females were only produced by 0×0 families and one 1×1 family, and were absent from any 2×2 DHS families. Zero spot males, however, were produced by all 0×0 families, all but one of the 1×1 families and all but one of the 2×2 families. Two spot females were produced by all but one (a 0×0) family, while 2 spot males were produced by all 2×2 families, but only two 1×1 families and one 0×0 family (Table 1). These results demonstrate that alleles are sufficiently segregating in our experimental design to perform heritability estimates.

**Table 1.** Dorsal hindwing eyespot number (DHEN) is heritable. Summary of DHEN data for offspring from 6 families each of 0×0, 1×1, and 2×2 DHEN crosses, separated by sex. Offspring with asymmetric DHEN are included in the average DHEN estimate.

|  | **0 x 0 DHEN Families** |  | **1 x 1 DHEN Families** |  | **2 x 2 DHEN Families** |  |
| --- | --- | --- | --- | --- | --- | --- |
|  | Females | Males | Females | Males | Females | Males |
| 0 DHEN | 46 | 159 | 2 | 47 | 0 | 14 |
| 1 DHEN | 55 | 15 | 45 | 36 | 19 | 38 |
| 2 DHEN | 24 | 2 | 43 | 9 | 67 | 27 |
| 3 DHEN | 2 | 0 | 4 | 0 | 6 | 0 |
| **Average DHEN** | **0.87** | **0.17** | **1.49** | **0.63** | **1.87** | **1.13** |

DHEN has a heritability of 0.4442 ± 0.264 for females, 0.5684 ± 0.306 for males, and 0.5029 ± 0.159 when the sexes were pooled. There was no significant interaction of parental phenotype and offspring sex on offspring phenotype (General linear Model with sex, parental phenotype, and sex*parental phenotype as parameters, AIC (Akaike Information Criterion) = 1498.204, effect tests: sex χ^2^=293.361, p<0.0001; parental phenotype χ^2^=271.56, p<0.0001; sex* parental phenotype χ^2^=0.032, p=0.8576).

### Genome-wide variation and linkage disequilibrium of the study population

To confirm the absence of substructure in our study population, we calculated measures of genetic variation and diversity between the samples displaying presence of DHEN (pre) and samples with absent DHEN (abs). As expected, the two groups showed very little genome-wide genetic divergence, with a genome-wide *F_ST_* equal to 0.0075. The absence of any population substructure between the two sampled phenotype groups was further demonstrated by complete overlap of the two groups in the PCA as well as little contribution of phenotype group to the observed variation in first and second Principal Components (Fig. 2A). Additionally, we observed very similar genome-wide nucleotide diversity in the group displaying presence of DHEN (pre, π = 0.0090) when compared to the group with absent DHEN (abs, π = 0.0083). Hence, we do not observe any apparent demographic substructuring of the study population that could potentially bias our genetic association analysis. Further, potential relatedness of individuals in our sampling was controlled for by the addition of a relatedness matrix in the generation of the linear mixed model used for testing association.

**Figure 2.**
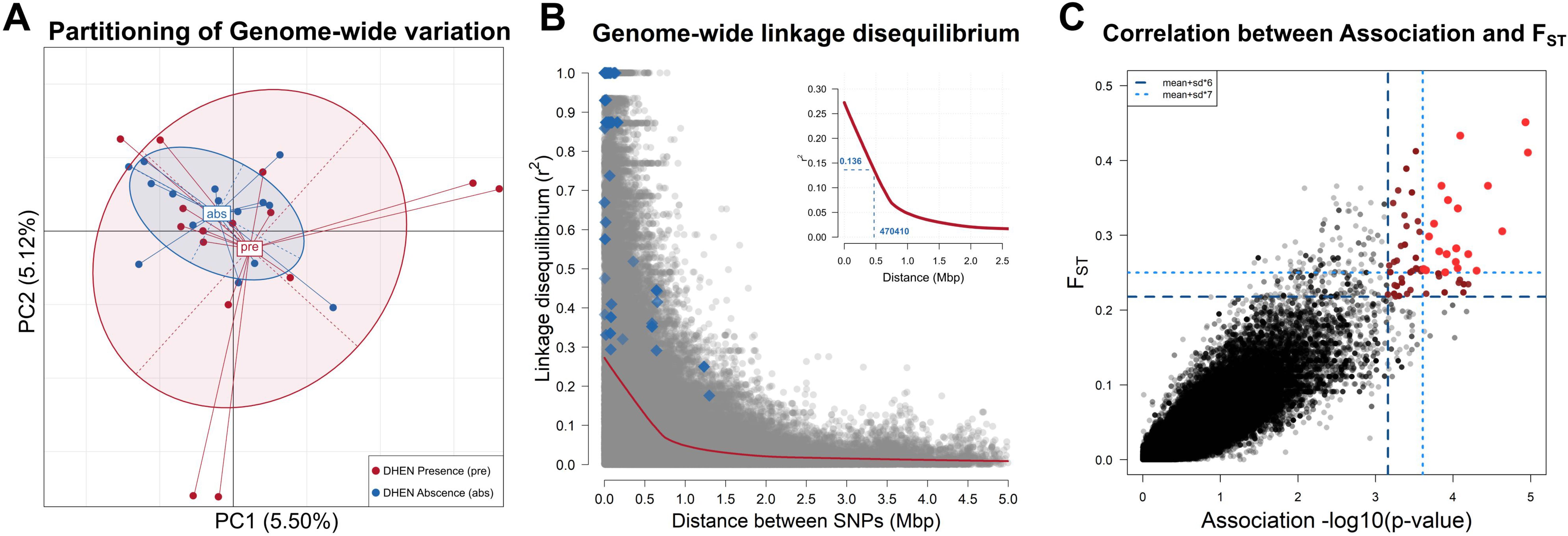
Suitability control for using the lab reared *B. anynana* population for GWAS. **(A.)** Principal Component Analysis (PCA) of the allelic variation observed in 5000 randomly-selected genome-wide SNPs in the study population across both phenotype groups, DHE presence (*pre*, red) and DHE absence (*abs*, blue). Ellipses display boundaries of the 95% confidence interval. Little contribution to variation in the principal components and overlap of the variation on both phenotype groups suggests lack of underlying demographic substructuring in the study population. **(B.)** Genome-wide linkage disequilibrium (LD) in the *B. anynana* study population. Grey dots represent LD values for a SNP pairwise comparison. Blue diamonds show LD values for SNP pairwise comparisons across outlier SNPs. In red, Loess regression smoothed curve representing LD decay. Insert: Zoomed-in LD decay curve, indicating distance at which LD is halved (470,410bp) and corresponding r^2^ value (0.137). **(C.)** Correlation between association (-log10(*p*-value)) and *F_ST_* between study groups. Dots denote the values for association and *F_ST_* for all SNPs in the dataset. Black dots represent non-outlier SNPs. Dark and light red dots represent outlier SNPs above the 6 and 7 standard deviation above the mean cutoff, respectively (dark and light blue dashed line).

After calculating genome-wide estimates of linkage disequilibrium decay (Fig. 2B), we observed a max smoothed r^2^ value of 0.272, and a halving of r^2^ within 470 kb (Fig. 2B). This window size suggests that average linkage blocks are around 500 Kb in length, and that variants within this distance may be in strong linkage disequilibrium.

### Association mapping of dorsal hindwing spot number variation

After mapping and genotyping RAD loci across the *B. anynana* reference genome, we identified a total of 216,338 SNPs shared between the two different genotyping strategies used (see methods). The distributions and correlations between the distributions of *F_ST_* and association are strongly positively correlated across all SNPs with a slight bias of association values toward more significant values for intermediate *p*-values, however, below our outlier thresholds (Figure 2C, Figure S1). We observed 99 SNPs on 25 different scaffolds with both *F_ST_* and association with DHEN variation at least 6 standard deviations above the mean *F_ST_* and association estimates. These SNPs span 28.5 Mb (6.01 %) of the reference genome assembly (Fig. 3). When using a more stringent threshold at 7 standard deviations, we observe a total of 28 outlier SNPs in 8 scaffolds, spanning 7.78 Mb (1.64%) of the genome. Ordering these scaffolds along the contiguously assembled *Heliconius melpomene* genome suggests that the 25 scaffolds with outlier SNPs above 6 standard deviations from the mean *F_ST_* and association belong to 21 different regions in the genome, whereas the smaller subset of 8 scaffolds that have outlier SNPs above 7 standard deviations belong to 6 different regions in the genome (Fig. 4).

**Figure 3.**
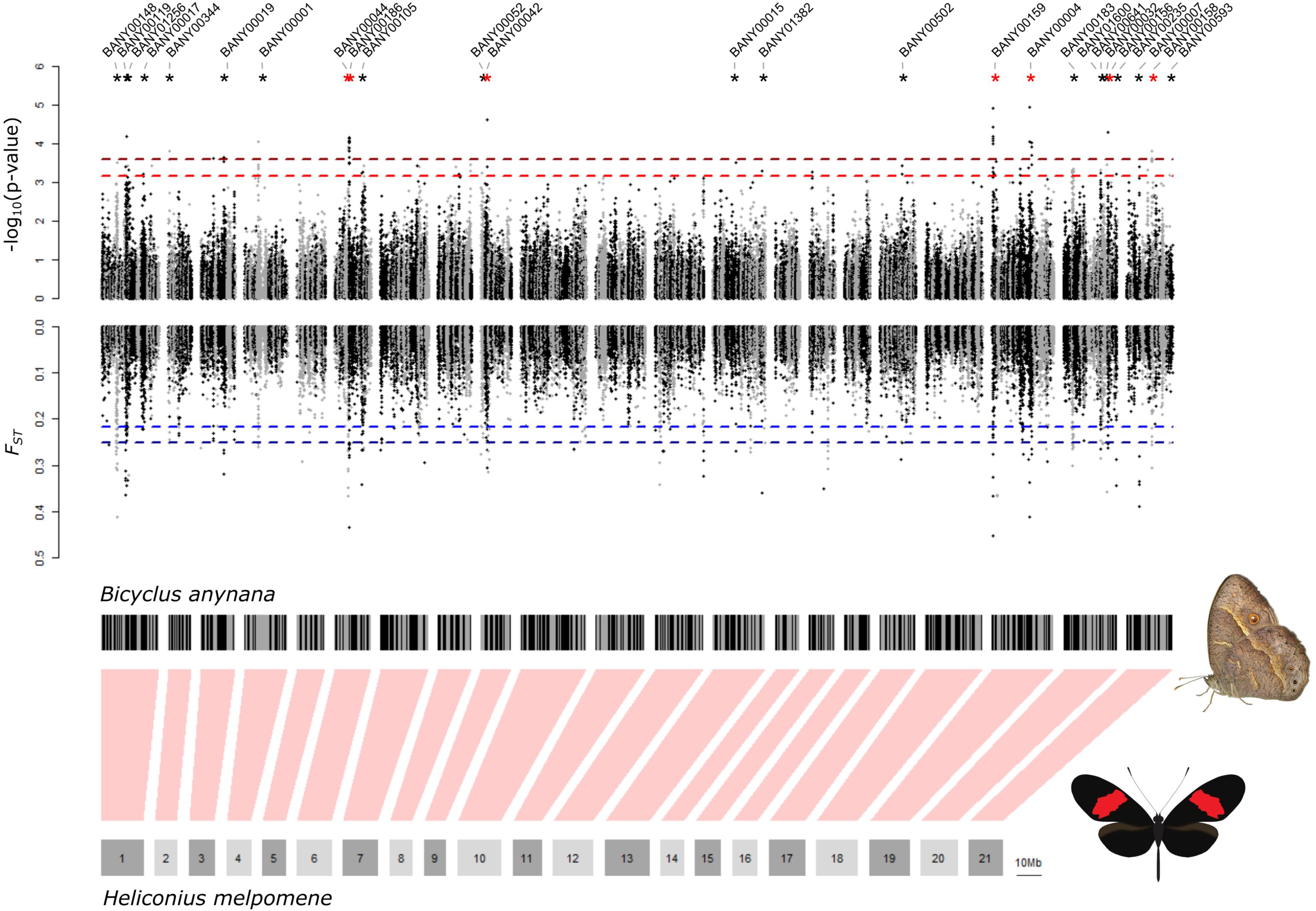
Genome-wide association with dorsal hindwing eyespot number. **(A.)** Plots show genomic association to dorsal hindwing eyespot number (top) and *F_ST_* between individuals with a different number of dorsal hindwing eyespots (bottom). Each dot represents a single SNP. Dashed lines represent the threshold for detecting a significant genome-wide association (top, in red) and *F_ST_* (bottom, in blue) above 6 and 7 standard deviations (SD) above the genome-wide mean. Scaffolds containing both significant association and *F_ST_* outliers are marked with asterisks (black for SD > 6 and red for SD > 7). **(B.)** Genomic scaffolds from the *Bicyclus anynana* BANY.1.2 genome are arranged along the 21 chromosomes of the *Heliconius melpomene* v2 assembly. For ordering the *B. anynana* scaffolds along the *H. melpomene* genome, only matches with a minimum percentage of identity of 90% and a minimum alignment length of 200 bp were used. If scaffolds matched multiple *H. melpomene* chromosomes, the scaffold was positioned along the chromosome to which it had the most matches. Using this strategy 76.7% of the *B. anynana* genome scaffolds were aligned to the *H. melpomene* genome.

**Figure 4.**
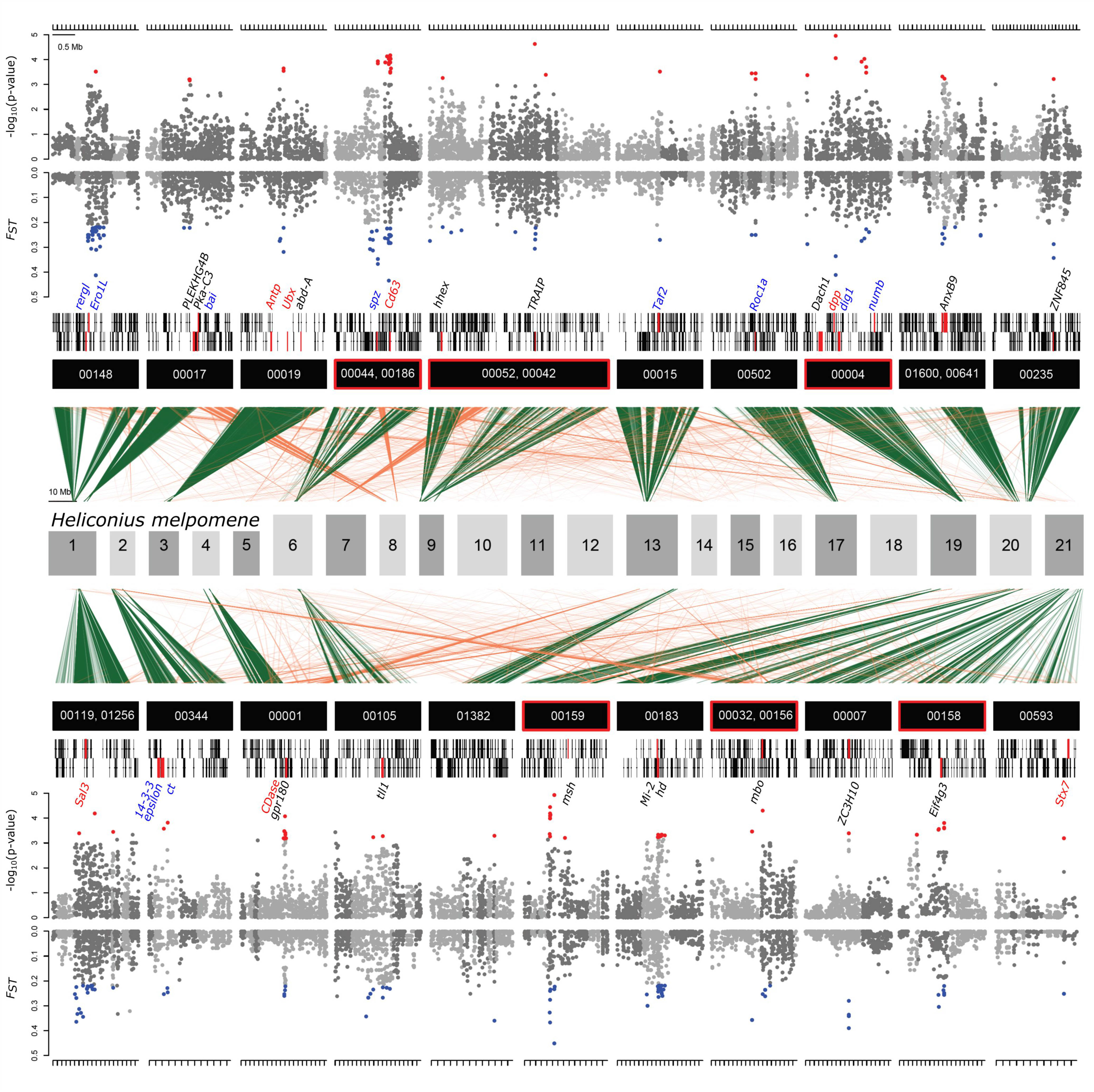
Zoom-in on putative genomic regions underlying dorsal hindwing eyespot number variation. Plots show genomic association to dorsal hindwing eyespot number (top) and *F_ST_* (bottom) between individuals with different dorsal hindwing eyespot numbers for scaffolds with significant outliers (red for association and blue for *F_ST_*). Each dot represents a single SNP. Light and dark gray dots correspond to alternating BANY.1.2 scaffolds mapping adjacently to the *H. melpomene* genome. Only names of scaffolds with significant outliers are listed. Scaffolds with SNPs above the SD 7 threshold are highlighted with a red box. Green and orange lines show matches of the *B. anynana* scaffolds (minimum percentage of identity of 70% and a minimum alignment length of 150 bp) to the *H. melpomene* v2 assembly. Green lines represent the most frequent matches of the scaffold to a *H. melpomene* chromosome, whereas orange lines represent matches to a different *H. melpomene* chromosome. Vertical black rectangles represent gene models with candidate genes highlighted in red. Names of genes that overlap or are close to an outlier SNP and that are reported in the literature to affect eyespot development are indicated in red. Names of genes that are within the same pathway as genes involved in eyespot development are indicated in blue. Names of other candidates are indicated in black.

### Candidate gene identification

Using the available Lepbase reference annotation for the *B. anynana* v1.2 assembly we identified the neighboring annotated genes and relative positioning of the 99 outlier SNPs on the 21 genomic regions. We observed a total of 742 annotated genes within our 500 kb windows around the outlier SNPs (Supplemental Table S2), of which 251 remained with the 28 outlier SNPs that are above the higher 7 standard deviation threshold. Only 25 of these 742 genes contained outlier SNPs in their introns or exons (Table 3, supplemental Table S2). On average, our outlier SNPs are 189 kb (±144 kb) from the nearest gene. This suggest that a significant portion of our outliers are intergenic and potentially associated with regulatory variants. Only a few SNPs mapped within a gene’s coding sequence, of which 9 resulted in synonymous and 2 in non-synonymous mutations (Table 3, Supplemental Table S2). Thus, in order to identify the most promising candidate genes that may be linked to the non-coding associated SNPs and regulating DHEN, we used our current knowledge of butterfly developmental genetics and identified 35 such genes (Table 2, Figure 4). Among those, 7 candidates were previously associated with eyespot development, i.e., *CDase, Dpp, Antp, Ubx, Sal3, Cd63, and Syntaxin 7*, and the remaining 28 are linked to eyespot number variation here for the first time (Table 2). The latter include genes related to Notch, Toll, and Ras signaling pathways, wing morphogenesis, dorso/ventral pattern formation, and pigment biosynthesis and a number of molecules with regulatory and signaling functions (Table 2).

**Table 2.**
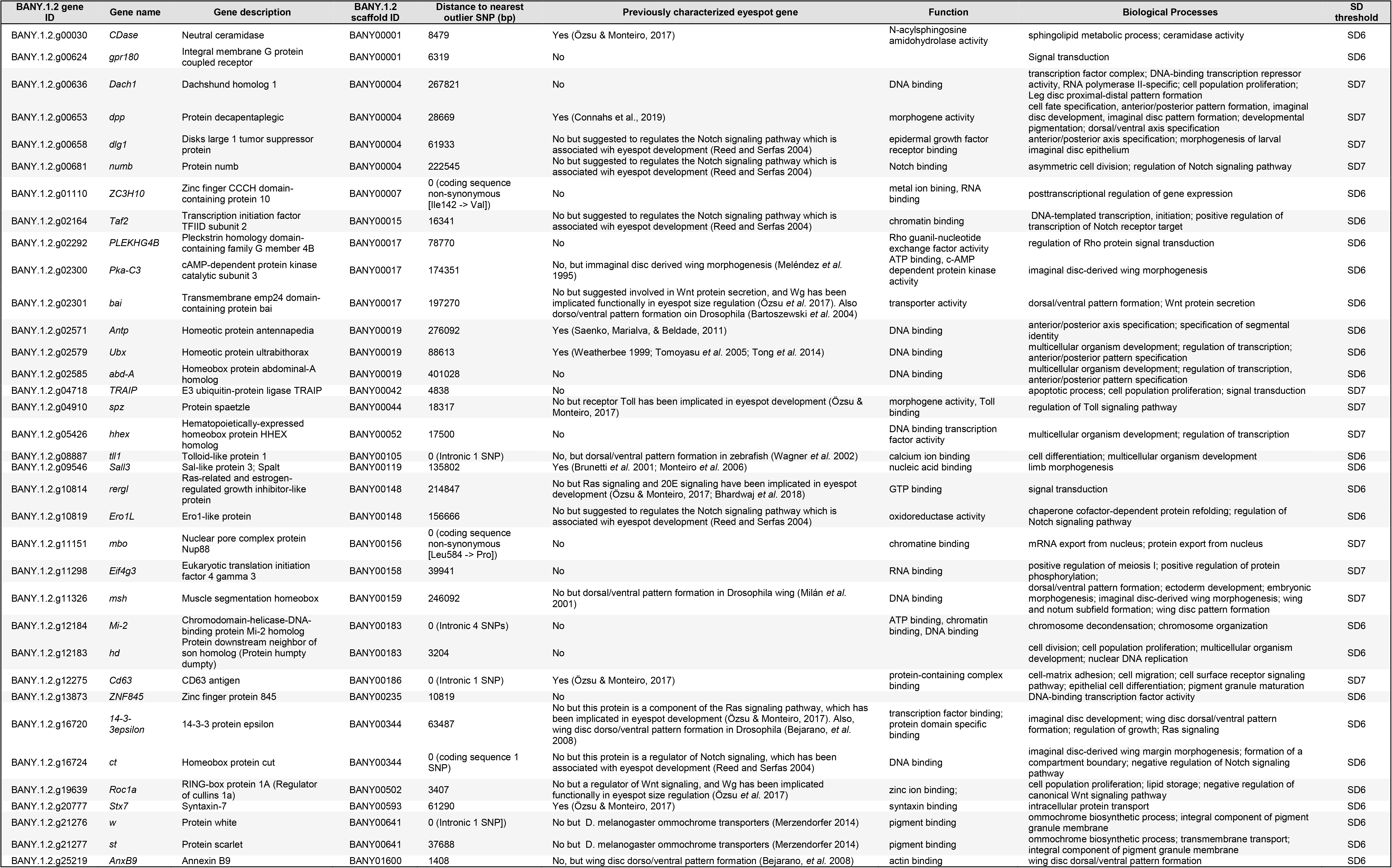
DHEN candidate genes. BANY.1.2 gene ID, gene name, gene description and BANY.1.2 scaffold ID all refer to the *B. anynana* v1.2 genome assembly and annotation (Nowell *et al.* 2017). Distance to nearest SNP refers to the distance of each of the candidate genes to one of our identified SNPs in bp. Genes with SNPs inside their sequence have additional information describing the SNP annotation. Eyespot regulatory networks describes direct or indirect evidence on the role of each of our identified candidates within the gene networks responsible for eyespot development, in addition to suggested functions within this network. Genes were identified based on two possible association thresholds, as described in methods.

**Table 3.**
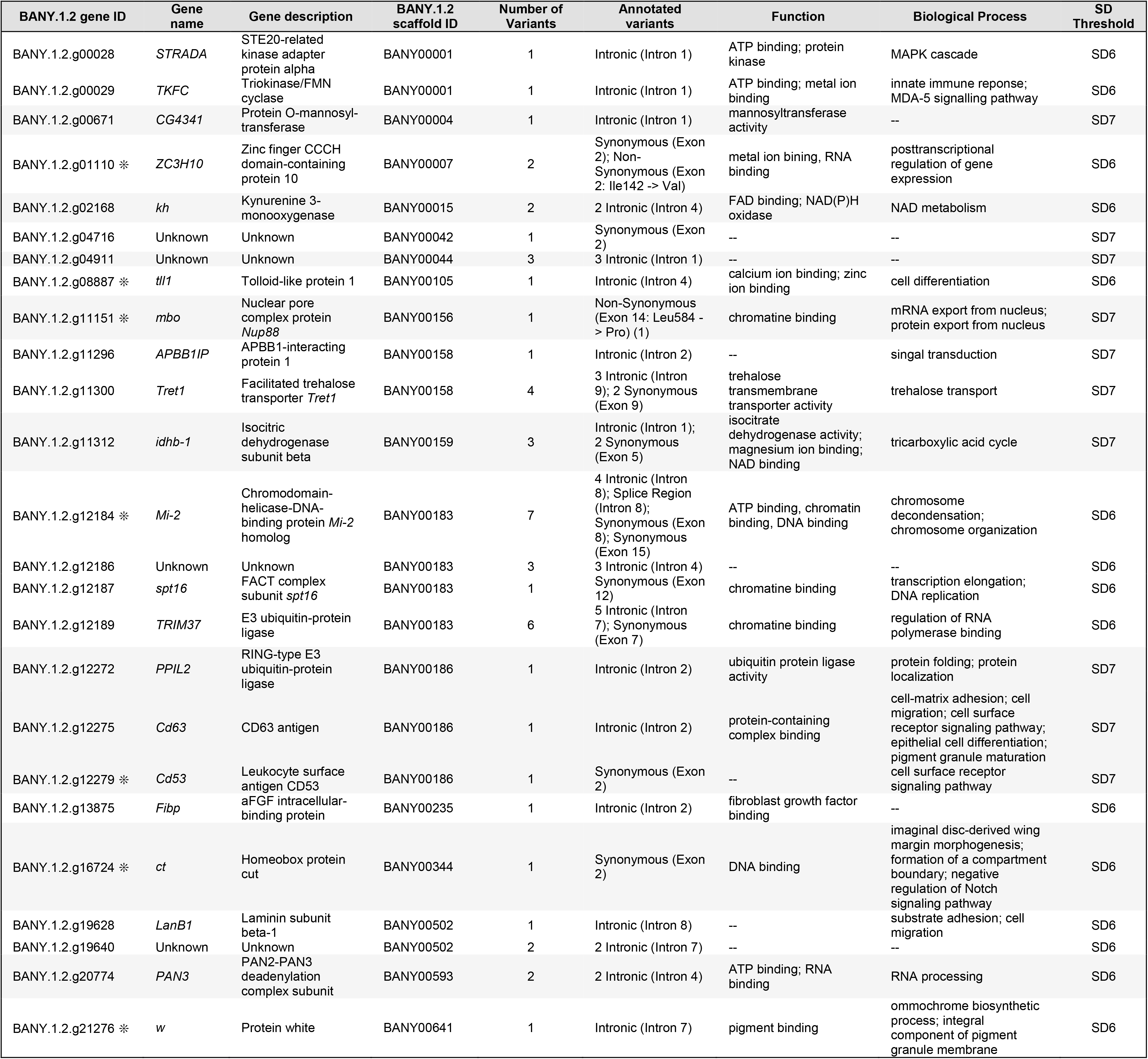
Genes containing outlier SNPs. BANY.1.2 gene ID, gene name, gene description and BANY.1.2 scaffold ID, and Function/Biological process all refer to the *B. anynana* v1.2 genome assembly and annotation (Nowell *et al.* 2017). Genes were identified based on two possible association thresholds, as described in methods. The number of variants refers to the number of outlier SNPs present inside the gene. Each individual variant is annotated, and its position within the gene’s sequence and impact on amino acids changes are described. Asterisks in gene IDs represents a gene that overlaps with our DHEN candidates in Table 2.

## Discussion

The use of population genomic analyses, including association mapping, GWAS, and *F_ST_* scans, has been extensively used in natural populations to identify the genomic signature underlying a number of biological processes such as hybridization, speciation, selection and ecological adaptation (Reviewed in Campbell, Poelstra, & Yoder, 2018; Narum, Buerkle, Davey, Miller, & Hohenlohe, 2013; Rellstab, Gugerli, Eckert, Hancock, & Holderegger, 2015). In our work, we applied a population genomics approach to characterize the genetic basis of dorsal hindwing eyespot number (DHEN) variation in the butterfly *B. anynana* via the comparison of genomic diversity between individuals from a single laboratory-maintained, freely breeding population, that display moderate to high narrow-sense DHEN heritability. Like natural populations, we demonstrated that the laboratory population maintains homogenization of the genetic variation across the genome due to a lack of demographic substructuring, and thus provides a powerful system to detect associations between SNPs and the trait of interest (DHEN variation).

The combined results of our heritability and genome-wide association study suggest that variation in dorsal hindwing eyespot number in *B. anynana* is a complex trait regulated by multiple loci. Combining a RAD sequencing genotyping approach with genome-wide population differentiation and association mapping, we identified up to 21 and 6 genomic regions associated with DHEN variation in *B. anynana*, at different levels of stringency (Figure 3 and 4). Within the 21 genomic regions, we observed a total of 742 annotated genes (Supplemental Table 2), which we narrowed down to 35 potential candidates (Table 2). Within the stronger associated 6 genomic regions (Figure 4), we reduced the list of potential candidate genes to a total of 11 (Table 2). Below we address the possible relationship of these candidates with eyespot development and DHEN variation.

### Known eyespot developmental genes associated to DHEN variation

Among our identified candidates, we observed a total of seven genes previously implicated in eyespot development in *B. anynana*: *CDase, Dpp, Antp, Ubx, Sal3, Cd63*, and *Syntaxin 7* (Brunetti *et al.* 2001; Tong *et al.* 2014; Özsu and Monteiro 2017; Connahs *et al.* 2019) (Table 2). From these, only *dpp* and *Cd63* remained associated at the highest significant threshold of 7 SD. Most of these genes are within or near the area of strongest association, and only *Cd63* has an associated SNP within an intron (Table 3 and Figure 4). Neutral Ceramidase (*CDase*; BANY.1.2.g00030) is an enzyme involved in the metabolism of sphingolipids which is down-regulated in eyespots relative to flanking wing tissue (Özsu and Monteiro 2017). Sphingolipids appear to function in a variety of cell signaling pathways but are still poorly understood (Hannun and Obeid 2017). Decapentaplegic (*Dpp*; BANY.1.2.g00653) is a known developmental gene expressed in dynamic patterns during eyespot center formation during the larval stage (Monteiro *et al.* 2006; Connahs *et al.* 2019). *Dpp* is likely involved in the differentiation of the eyespot centers in *B. anynana* via a reaction-diffusion mechanism (Connahs *et al.* 2019). *Dpp* mRNA is expressed in regions around the developing eyespots in mid to late larval wings (Connahs *et al.* 2019) in anti-colocalized patterns to Armadillo, a transcription factor effector of the Wingless signaling pathway. Armadillo is expressed in the actual eyespot centers in late larval stages (Connahs *et al.* 2019), as is Antennapedia (*Antp*; BANY.1.2.g02571), a hox gene (Saenko *et al.* 2011; Tong *et al.* 2014; Özsu and Monteiro 2017). Antp is also involved in the differentiation of the thoracic limbs in *Bombyx* moths (Chen *et al.* 2013). Ultrabithorax (*Ubx*; BANY.1.2.g02579) is another hox gene that gives insect hindwings a different identity from forewings (Weatherbee *et al.* 1999; Tong *et al.* 2014). In *B. anynana*, this gene is expressed across the whole hindwing, a conserved expression pattern observed across insects, but has additional, stronger, expression in the eyespot centers of the hindwing only, something that is not seen in other butterflies with eyespots such as *Junonia coenia* (Tong *et al.* 2014). Over-expression of *Ubx* led to eyespot size reductions in both fore and hindwings of *B. anynana* (Tong *et al.* 2014) whereas absence of *Ubx* in clones of cells in the hindwing of *J. coenia* (via a spontaneous unknown mutation) (Weatherbee *et al.* 1999) led to eyespot enlargements (of Cu1 hindwing eyespots to sizes that match Cu1 forewing eyespot sizes). These results suggest a repressing role of *Ubx* for eyespots on the hindwing. Genetic variation at *Ubx* might contribute to eyespot number variation via a threshold-like mechanism acting on eyespot size. Sal-like protein 3, SALL3, also known as *Spalt* (*Sal*; BANY.1.2.g09546) is a transcription factor that is expressed in eyespot and spot centers in a variety of butterflies including *B. anynana* (Brunetti *et al.* 2001; Oliver *et al.* 2012; Shirai *et al.* 2012) and was recently shown to be required for eyespot development in both *V. cardui* and *J. coenia* butterflies (Zhang and Reed 2016). The trespanin Cd63 (Cd63; BANY.1.2.g12275) is a protein associated with exosomes, which in turn have been associated with wingless signaling in *Drosophila* (Beckett *et al.* 2013). This gene is the only candidate previously characterized as up-regulated in eyespots (Özsu and Monteiro 2017) to contain an association within its gene module, and more specifically a SNP in its 2nd intron. Lastly, Syntaxin 7 (*Stx7*; BANY.1.2.g20777) has a similar function to *Syntaxin* 1a, a gene that is down-regulated in eyespots (Özsu and Monteiro 2017). Genetic variation either linked with the protein coding sequence of these genes, or more likely with their regulatory region is likely affecting eyespot number variation in hindwings.

### Novel candidate genes associated to DHEN variation

In addition to the above eyespot-associated genes, we also identified a group of 28 candidates that have not been associated with eyespot development before. However, many of these genes are part of signaling pathways previously associated with eyespot development such as the Notch, Ras, and Toll signaling pathways (Table 2) (Özsu and Monteiro 2017), and others with roles in wing morphogenesis, dorso/ventral pattern formation, and pigment biosynthesis.

A total of five genes (*dlg1, numb, Taf2, Ero1L*, and *ct*) implicated in our study (Table 2) are known to interact with the *Notch* eyespot-associated gene (Reed and Serfas 2004). Disks large 1 tumor suppressor, (*dlg1*; BANY.1.2.g00658), numb (*numb*; BANY.1.2.g00681), Ero1-like protein, (*Ero1L*; BANY.1.2.g10819) and ct (*ct*; BANY.1.2.g16724) are known to interact and regulate the *Notch* (*N*) signaling pathway in D. melanogaster (Cheah *et al.* 2000; Tien *et al.* 2008; Li *et al.* 2009b). The gene *ct* is the only one of these genes with an associated SNP, causing a synonymous mutation. The existence and role of these interactions are unknown in *B. anynana*, as is the role of the *Notch* receptor itself. However, the *Notch* receptor has a dynamic pattern of expression (Reed and Serfas 2004) that is very similar to that of *Distal-less*, a gene that has recently been implicated in setting up the eyespot centers likely via a reaction-diffusion mechanism (Connahs *et al.* 2019). Genetic variation at these three genes could be interacting with the eyespot differentiation process through unknown mechanisms.

We also identified new members of the Ras and Toll signaling pathways that have previously been associated with eyespot development (Özsu and Monteiro 2017). 14-3-3 epsilon (BANY.1.2.g16720) is a member of the Ras signaling pathway (Chang and Rubin 1997) and Ras-related and estrogen-regulated growth inhibitor-like protein (*rergl*; BANY.1.2.g10814), is a Ras-related superfamily gene with unique characteristics (Finlin *et al.* 2001). Protein spaetzle (*spz*; BANY.1.2.g04910) is a ligand that enables the activation of the Toll pathway in *Drosophila* (Yamamoto-Hino and Goto 2016). The role of *spz* is currently unknown in the context of eyespot development but this ligand could be an interesting target of further study. Our data suggests that genetic variation at these loci might also be regulating hindwing eyespot number variation.

Interestingly, some of our candidate genes involve *Drosophila* embryonic, wing-specific and dorso/ventral patterning genes. The latter function is in line with eyespot number variation being observed on the dorsal but not on the ventral surface of the hindwing. cAMP-dependent protein kinase catalytic subunit 3 (*Pka-C3*; BANY.1.2.g02300), Transmembrane emp24 domain-containing protein baiser (*bai*; BANY.1.2.g02301), Muscle segmentation homeobox (*msh*; BANY.1.2.g11326) and RING-box protein 1A (*Roc1a*; BANY.1.2.g19639) are all involved in embryonic development or wing morphogenesis in *Drosophila*. *Pka-C3* is specifically expressed in wing disc and leg discs of third instar larvae (Meléndez *et al.* 1995). *bai* is involved in the regulation of a receptor of *dpp* in *Drosophila* embryos (Bartoszewski *et al.* 2004) and in *Wnt* protein secretion (Li *et al.* 2015), both key signaling pathways implicated in eyespot size regulation (Özsu and Monteiro 2017; Connahs *et al.* 2019). *Msh* specifies dorsal cell fate in the *Drosophila* wing (Milán *et al.* 2001), and *Roc1a* negatively regulates the canonical Wnt signaling (Roberts *et al.* 2012), a key pathway for eyespot development (Özsu *et al.* 2017). Lastly, AnxB9 has been proposed to be involved in wing morphogenesis and wing dorsal pattern identity in *Drosophila* (Bejarano *et al.* 2008).

Among our candidates we also have two transporters of pigment precursors into pigment cells responsible for a fruit fly’s eye color: White (*w*; BANY.1.2.g21276) and scarlet (*st*; BANY.1.2.g21277). In our study *white* is one of the few genes that has an associated SNP in the 7th intron. In *Drosophila* these two genes are required, together with *Brown*, for normal pigmentation of the compound eye (Merzendorfer 2014). The two transporters white and scarlet have been suggested to import precursors of ommochrome pigments into butterfly scales, which are necessary for yellow, red and brown colorations (Hines *et al.* 2012). It is unclear, however, how these genes might play a role in the regulation of eyespot number variation.

Finally, we have identified several candidate genes with roles in pre- and post-transcriptional regulation (*Dach1, ZC3H10, abd-A, hhex, tll1, mbo, Eif4g3*, and *Mi-2*) and signal transduction (*gpr180, PLEKHG4B, and TRAIP*) that might play a role in DHEN variation. Of particular interest is Dachshund homolog 1 (*Dach1*; BANY.1.2.g00636), a gene that is involved in limb development and that is activated by Distal-less in *Drosophila* (Estella *et al.* 2012). Another is abd-A (abdominal-A; BANY.1.2.g02585), one of only two genes providing butterfly’s hindwing chromatin regulatory identity (Lewis and Reed 2019). Other genes (*ZC3H10, tll1, mbo*, and *Mi-2*) have SNPs within introns or SNPs leading to non-synonymous mutations that can possibly provide them new binding affinity (Table 3). One of these intronic variants is annotated as putatively affecting a splice region within an intron in Chromodomain-helicase-DNA-binding protein *Mi-2* homolog (*Mi-2*; BANY.1.2.g12184). Moreover, we observed non-synonymous mutations in two different genes: Nuclear pore complex protein Nup88 (*mbo*; BANY.1.2.g11151) with a Leucine to Proline substitution at position 584, and Zinc finger CCCH domain-containing protein 10 (*ZC3H10*; BANY.1.2.g01110) with a Isoleucine to Valine substitution at position 142 (Table 3; Supplemental Table S2). Future functional studies will be required to further implicate these genes in eyespot development and eyespot number regulation.

### Effects of non-coding mutations in the evolution of eyespot number variation

After identifying 21 regions of the *B. anynana* genome associated with DHEN variation and characterizing the relationship of identified SNPs with nearby genes, we observed that the majority of SNPs (92.7%) fall outside coding sequences. Only two genes, *mbo* and *ZC3H10* contain variants annotated as non-synonymous mutations of unknown effect on the resulting protein, while a third, *Mi-2* possesses an annotated splice region variant. Overall, DNA variation at non-coding loci suggests a complex *cis*-regulatory landscape controlling DHEN throughout the mediated expression of the nearby genes described above. Indeed, *cis*-regulatory elements are thought to have profound implications in the evolution of morphological diversity (Carroll 2008). Particularly, they have been associated with variation in pigmentation patterns in a wide variety of animal systems, including the evolution of eggspot pigmentation patterns in cichlids (Santos *et al.* 2014), wing pigmentation patterns in *Drosophila* (Werner *et al.* 2010; Koshikawa *et al.* 2015), divergent pigmentation patterns in capuchino seedeater finches (Campagna *et al.* 2017), and variation in red, black and yellow color patterns in *Heliconius* butterflies due to regulatory changes in the *optix, WntA, cortex*, and *aristaless* genes (Reed *et al.* 2011; Supple *et al.* 2013; Martin and Reed 2014; Van Belleghem *et al.* 2017; Westerman *et al.* 2018) among others. In the case of eyespot number variation, regulatory mutations around the genes identified here might disrupt the reaction-diffusion mechanism of eyespot center differentiation (Connahs *et al.* 2019), or later processes of eyespot center signaling, that eventually translate to presence or absence of an eyespot in particular wing sectors.

The genetic variation uncovered in this work affects eyespot number variation on the dorsal surface but not on the ventral surface of the wing. Thus, our work suggests that the genetic variants identified with our analysis affect eyespot number in a surface-specific manner. This surface-specific regulation is potentially mediated via *apA*, a previously identified dorsal eyespot repressor (Prakash and Monteiro 2018). The polygenic nature of our results proposes that genetic variation at the loci identified above, e.g., *Antp, Ubx, dpp*, etc, rather than at the *apA* locus itself regulates dorsal eyespot number. We can speculate that changes in gene expression at the identified loci might impact the expression of *apA* in specific wing sectors on the dorsal surface, ultimately controlling eyespot differentiation in those regions.

Finally, the use of a genome-wide sequencing strategy allowed us to discover a series of independent loci that appear to contribute to DHEN in *B. anynana*. These loci, predominantly composed of (or linked to) polymorphisms in non-coding DNA, suggest that changes in DHEN are mostly occurring in regions that regulate the expression of previously known eyespot-associated genes. This study identifies a polygenic architecture of hindwing eyespot number variation that might be most likely mediated by epistatic interactions among a set of developmental genes. Thus, while our work has enriched the list of genes involved in eyespot number variation, it also confirms that variation at multiple genes, rather than at a single top master regulator or input-output gene (such as *shavenbaby* or *achaete-scute*) is involved in regulating number of serial homologs. This highlights a more complex, but still poorly understood, genetic architecture for serial homolog number regulation.

## Acknowledgements

We thank Elizabeth Schyling for help rearing families for heritability analysis, and Robert Rak and Chris Bolick for rearing corn plants for the larvae. We also thank the UPR-RP Sequencing and Genomics Facility and the UPR-RP High Performance Computing Facility for additional support in library preparation and computational analysis. This work was funded by a NSF IOS-110382 Doctoral Dissertation Improvement Grant to ELW and AM, a NSF PR-LSAMP Bridge to the Doctorate Program (NSF Grant Award HRD1139888) to AGRC, NSF grant IOS-1656389 to RP, NSF grant OIA 1736026 to BC and RP, and a Puerto Rico INBRE Grant P20 GM103475 from the National Institute for General Medical Sciences (NIGMS), a component of the National Institutes of Health (NIH) and Ministry of Education, and Singapore grants (MOE R-154-000-602-112 and MOE2015-T2-2-159) to AM.

